# Detection and antibiotic susceptibility of pathogenic *Escherichia coli* isolated from the final effluent of two wastewater treatment Plants in the Eastern Cape Province, South Africa

**DOI:** 10.1101/160697

**Authors:** Osuolale Olayinka, Okoh Anthony

## Abstract

Wastewater is an important reservoir for *Escherichia coli* and can present significant acute toxicity if released into receiving water sources without being adequately treated. To analyze whether pathogenic *E. coli* strains that cause infections are in treated effluent and to recognize antibiotic profile. 476 confirmed isolates from two treatment Plants were characterized for the presence of various *E. coli* pathotypes. A total of 8 pathotypes were screened and only four were confirmed. UPEC was about 5.7% followed by EAEC at 2.3%, NMEC at 1.1% and EPEC at 0.6%. Antibiotic susceptibility patterns of *E. coli* pathotypes such as UPEC showed low resistance to antibiotics like meropenem (100%), cefotaxime (100%) and gentamicin (88.9%). The pathotype also showed high degrees of resistance to tetracycline (74.1%), ampicillin (74.1%) and cephalothin (66.7%). Other *E. coli* pathotypes, EAEC, NMEC and EPEC, showed high sensitivity (100%) to meropenem, gentamicin and cefotaxime, and varying degree of resistances to ampicillin, tetracycline and cephalothin. The results of this study reveal that the two Plants discharge effluents with pathogenic *E. coli* and are reservoir for the bacteria into receiving water sources. In summary, this finding raises the possibility that at least some pathogenic *E. coli* pathotypes are getting into the environment through WWTPs and represent potential route for enteropathogenic infection. In addition, certain pathotypes may have acquired resistance properties, becoming a potential cause of drug resistance infection. This study reveals inadequacy of the plants studied to produce effluents of acceptable quality.

## Introduction

Wastewater treatment Plants are important for managing and treating polluted used water. The management of this wastewater is crucial to averting environmental pollution, which could endanger public health (West and Mangiameli, 2000). The conventional system of wastewater treatment reduces the quantity of enteric bacteria. However, poorly treated wastewater in any of the treatment processes could hinder the effectiveness of any disinfectant applied to deactivate these organisms (Anastasi *et al*., 2012).

The incomplete removal of pathogens and antibiotic-resistant bacteria from wastewater has consequently introduced treated but contaminated wastewater effluents into natural water resources, escalating the risk of infection (Dolejska *et al*., 2011). *E. coli* with virulence characteristics of uropathogenic strains was reported to survive the treatment processes of sewage treatment Plants (STPs) and also found to be present in environmental water receiving effluent discharges from STPs (Anastasi *et al*., 2010, 2012).

Aquatic environments are natural reservoirs of antibiotic-resistant bacteria, and wastewater treatment Plants (WWTPs) are among the primary water reservoirs of these microorganisms. Antibiotic resistance genes in bacteria in water environments are a global concern and have increased dramatically in the recent years. Broad range of antibiotics resistance encoding genes from microorganisms have been found in wastewater effluents, surface water, river water, groundwater and drinking water (Dolejska *et al*., 2011). Multi-drug resistance has been shown in *E. coli* (Shariff *et al*., 2013).

To this extent, studies from several provinces in South Africa on wastewater effluents and water bodies have demonstrated the presence of pathogenic and antibiotic resistance *E. coli* (Omar and Barnard, 2010; Olaniran *et al*., 2009; Phokela *et al*., 2011).

The present study is a follow up study by (Osuolale and Okoh, 2015a, 2017, 2015b) undertaken to assess the quality of treated effluent discharged from wastewater treatment Plants in Eastern Cape, South Africa. This study was done as part of a wider study that included three other WWTPs, though those sites were used for viral sampling rather than bacterial sampling (Osuolale and Okoh, 2017) The discharge to water bodies was tested for pathogenic *E. coli* and their antibiotic profiles. The areas of study are unique in their semi-rural and semi-urban features. Our study hopes to provide insights into the presence of pathogenic *E. coli* in treated effluent.

### Materials and Methods

#### Study area and sampling procedure

The Plant A wastewater treatment works is located at geographical location of longitude 33° 00’ 59”S and latitude 27° 51’ 48”E. The Plant is medium sized, with treatment capacity of 5Ml/day. The Bio-filter/PETRO (pond-enhanced treatment and operation) process treatment system is employed for the treatment of influent (DWAF, 2009) and the final effluent is discharged into the Umzonyana stream. The Plant B WWTP is located at geographical coordinate of Long. 27°23’47” S and Lat. 32°85’36” E. The Plant receives municipal domestic sewage and run-off water. The wastewater treatment Plant is medium size and an activated sludge system with design capacity of about 8 ML/day (DWAF, 2009). The Plant treats an average dry weather flow of 7000 m^3^/day and an average wet weather flow of 21 000 m^3^/day. The final effluent is discharged into the Mdizeni stream, which is a tributary of the Keiskamma River.

Samples were collected on a monthly basis from the final treated effluent (FE) for a period of 12 months (September 2012 to August 2013). Samples were collected in sterile 1.7 litre Nalgene bottles. 10% sodium thiosulphate was added to sampling bottles to neutralize the chlorine effect on the target organisms. Samples were stored and transported in chiller boxes to the Applied and Environmental Microbiology Research Group (AEMREG) laboratory at the University of Fort Hare, Alice, South Africa for analysis. The collected samples were processed within six hours. The sampling frequency and number of samples are as recommended in the Quality of Domestic Water Supplies Volume 2: Sampling Guide (DWAF *et al*., 2000).

Bacteriological analysis of the effluent samples for isolation was determined by membrane filtration according to (Sans, 2011). *E. coli* coliforms chromogenic Agar (Conda, Madrid) was used for the isolation of *E. coli*. It differentiates *E. coli* from the rest of the Enterobacteriaceae. *E. coli* is easily distinguishable due to the dark blue-greenish colony colour. The filters were placed on the agar and incubated at 37 °C for 24 hours. This was done in triplicate. The target colonies were counted and reported as CFU/100 ml. After 24 hrs incubation, counts in the suitable range (0-300 colonies) were recorded using manual counting and the results per dilution plate count were recorded.

### Genotypic identification of E. coli

#### Isolation of genomic DNA

Purified presumptive *E. coli* isolates were grown in Lubria broth (LB) overnight for crude DNA extraction. The ZR Fungal/Bacterial DNA MiniPrep by Zymo Research was used to extract genomic DNA following the manufacturer’s instructions; genomic extract was immediately used in the molecular identification of the isolated organisms. Alternatively, prior to the PCR reaction, the DNA extract was stored at -20 °C.

#### PCR

Primers specific for the confirmation of the *E. coli* isolates were used in the polymerase chain reaction. The primers specific for the *uid*A gene in *E. coli* previously developed and examined for specificity to faecal pollution were used in the molecular identification of the isolates. Molecular identification was done targeting the *uid*A gene using the forward (5’-AAAACGGCAAGAAAAAGCAG-3’) and reverse (5’ACGCGTGGTTAACAGTCTTGCG-3’) primers with a 147 bp expected amplicon (Dungeni *et al*., 2010). The reaction parameters were 94 °C for 2 min, 30 cycles of 94 °C for 1min, 62.7 °C for 90 min, and 72 °C for 1 min and a final extension at 72 °C for 5 min. The primers specific for pathotypes are shown in Table 1. PCR amplification was performed with a MyCycler thermal cycler PCR (BioRad). The PCR solution contained 2 × PCR mastermix, 100uM each of 1ul each of the forward and reverse primers. The total volume for the PCR reaction was 25µl, 5µl of template DNA from each bacterial strain was added to make the final 25µl reaction volume. Gel electrophoresis was performed on the PCR product and run on a 2% w/v agarose gel at 100 V for approximately 90 mins. The gel image was captured digitally and analyzed using the Uvitec, Alliance 4.7. The chromosomal DNA of the positive control was used as reference control for primer accuracy and specificity.

**Table 1:**
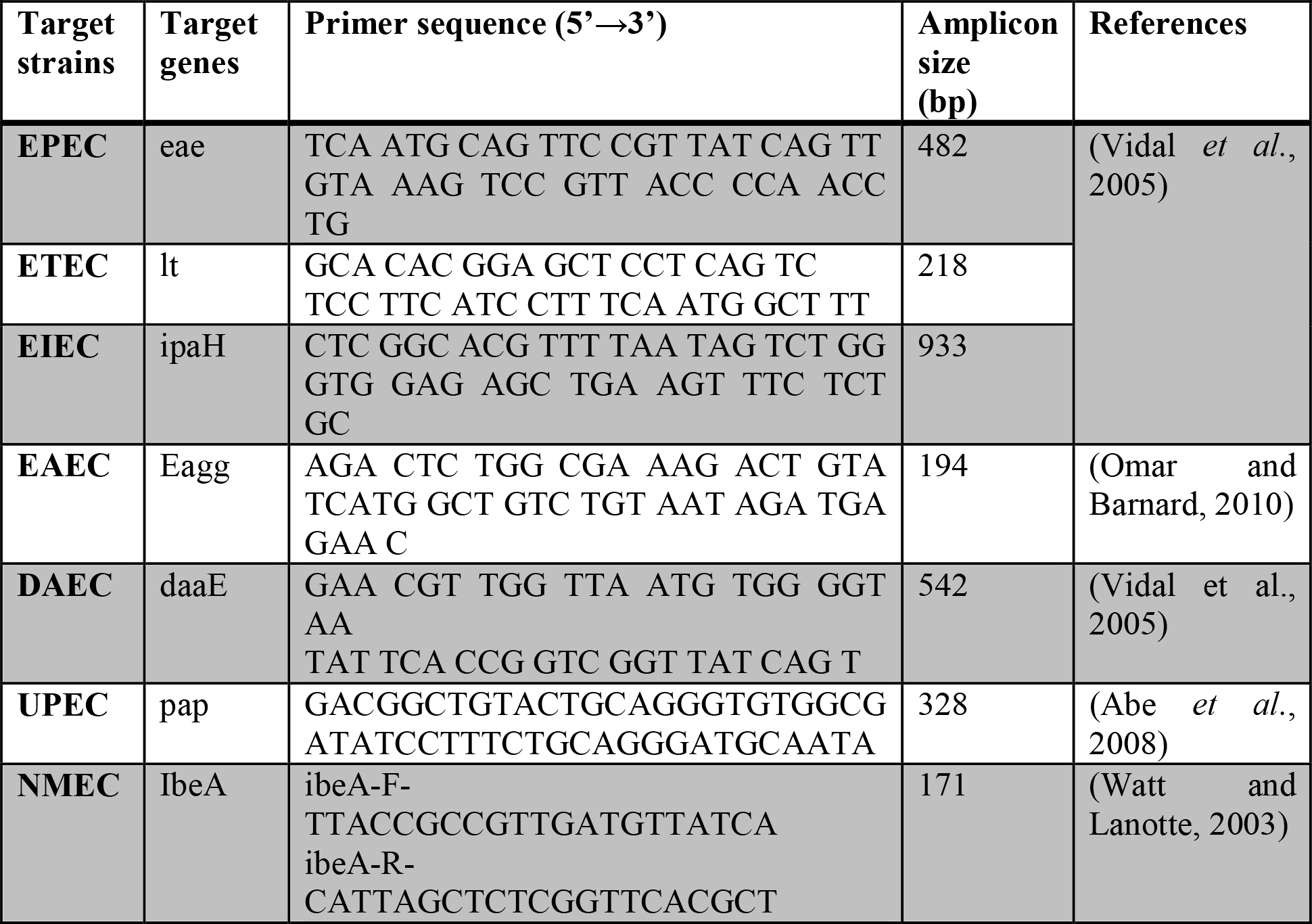
Primer pairs, expected amplicon size for characterization of *E. coli* pathotypes.

### Antimicrobial susceptibility testing

Antimicrobial susceptibility testing was done using the standard disc diffusion method on Mueller-Hinton agar (MH) (Conda, Madrid) as recommended by the Clinical and Laboratory Standards Institute (CLSI, 2012b). Fresh colonies (about 18 hrs old) from nutrient agar culture plates were picked into test tubes containing 5 ml sterile normal saline. The turbidity of the suspension was adjusted to 0.5 McFarland standards. Sterile swabs with bacterial suspensions were used to inoculate the MH agar plates by spreading uniformly on the surface of the agar. Selection of antimicrobials are based on the type of organism being tested and source of the isolates (CLSI, 2012b). Also, the antibiotics were selected as representatives of different classes of antibacterial drugs, to better depict the behaviour of the examined strains against these molecules. The antimicrobial susceptibility test for *E. coli* isolates was determined using the following antibiotic discs: ampicillin (10 µg), cefotaxime (30 µg), gentamicin (10 µg), meropenem (10 µg), tetracycline (30 µg), and cephalothin (30 µg) (Davies Diagnostics, SA) (CLSI, 2012b).

## Results

At Plant A, a total of 406 presumptive *E. coli* were isolated and 437 isolates were collected from Plant B (Table 2). During the study period, a total of 476 *E. coli* isolates from both Plants together were confirmed (Figure 1). About 5.7% (27) of the confirmed *E. coli* isolates were UPEC. The Plant A WWTP accounted for 77.8% (21) of the total UPEC isolates and Plant B accounted for 22.2% (6) (Table 3). Figure 2 (below) shows the PCR confirmation of the pap gene for UPEC. EAEC was the next most detected, accounting for 2.3% (11) of the total confirmed *E. coli* isolates (Figure 3). Plant A accounts for 81.8% (9) of the total confirm EAEC isolates, with 18.2% (2) at Plant B. Other confirmed pathotypes are NMEC (Figure 4), which was only detected in Plant A, and EPEC was only detected at Plant B. The other *E. coli* pathotypes like ETEC, EIEC and Diffuse-adhering *E. coli* were not detected at either Plant. The results of the *E. coli* pathotyping are as shown in Table 3, below.

**Table 2:**
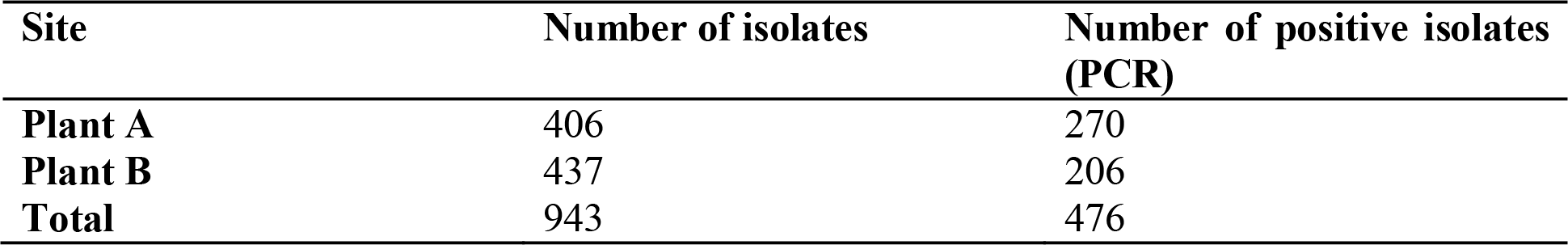
*E. coli* confirmation of the presumptive isolates.

**Figure 1:**
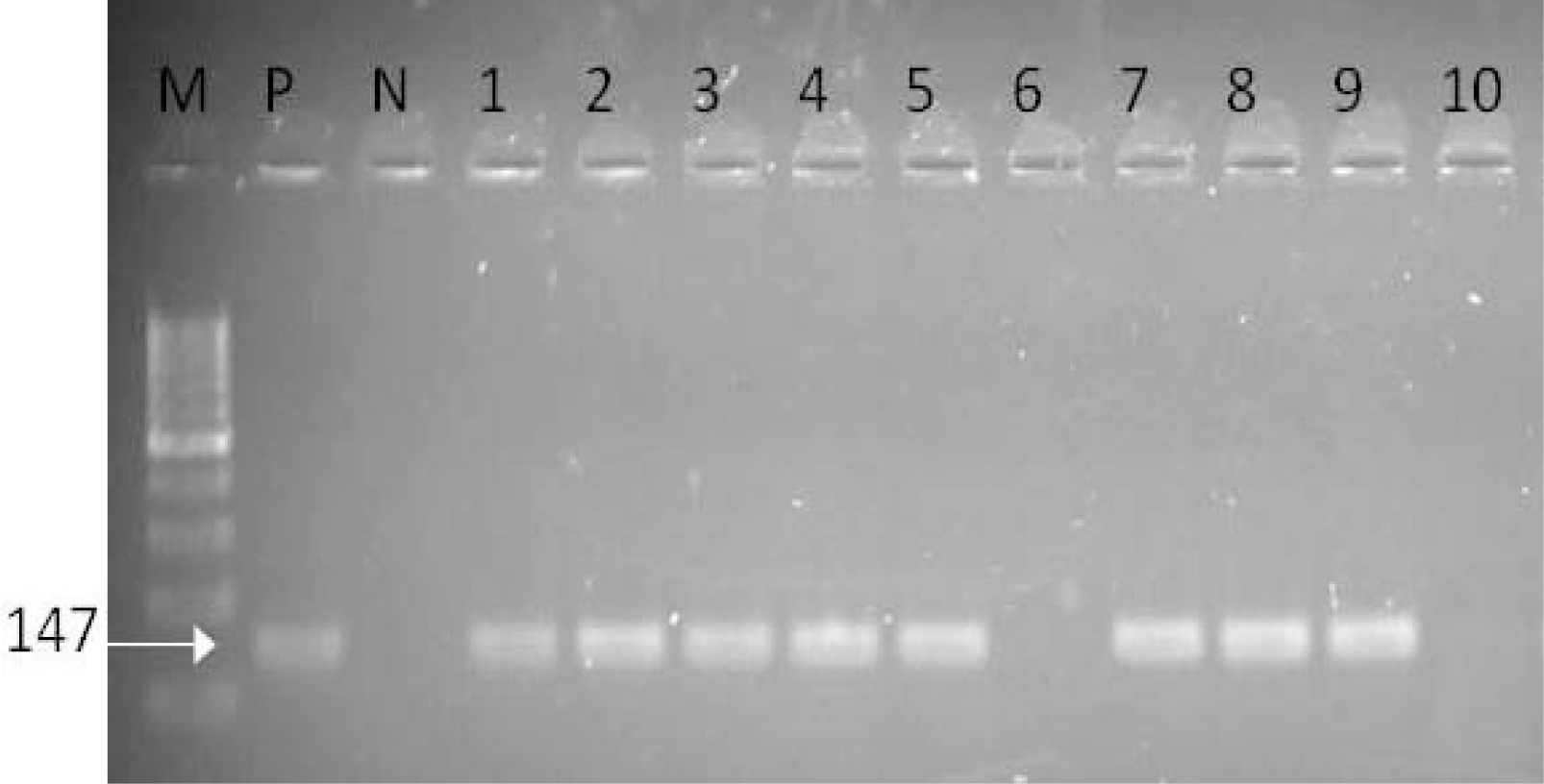
Agarose gel electrophoresis of uidA gene amplification products of *E. coli*. M: Molecular weight marker (100bp), P: *Escherichia coli* ATCC 8973 (Positive control), N: Negative control; Lanes 1-10: *E. coli* isolates

**Table 3:** Result of *E. coli* pathotyping.

| Pathotypes | Plant A (n = 270) | Plant B (n=206) |
| --- | --- | --- |
| Enteropathogenic *E. coli* (EPEC) | 0 | 3 |
| Enterotoxigenic *E. coli* (ETEC) | 0 | 0 |
| Enteroinvasive *E. coli* (EIEC) | 0 | 0 |
| Enteraggregative *E. coli* (EAEC) | 9 | 2 |
| Neonatal meningitis-associated *E. coli* (NMEC) | 5 | 0 |
| Uropathogenic *E. coli* (UPEC) | 21 | 6 |
| Diffuse-adhering *Escherichia coli* | 0 | 0 |

**Figure 2:**
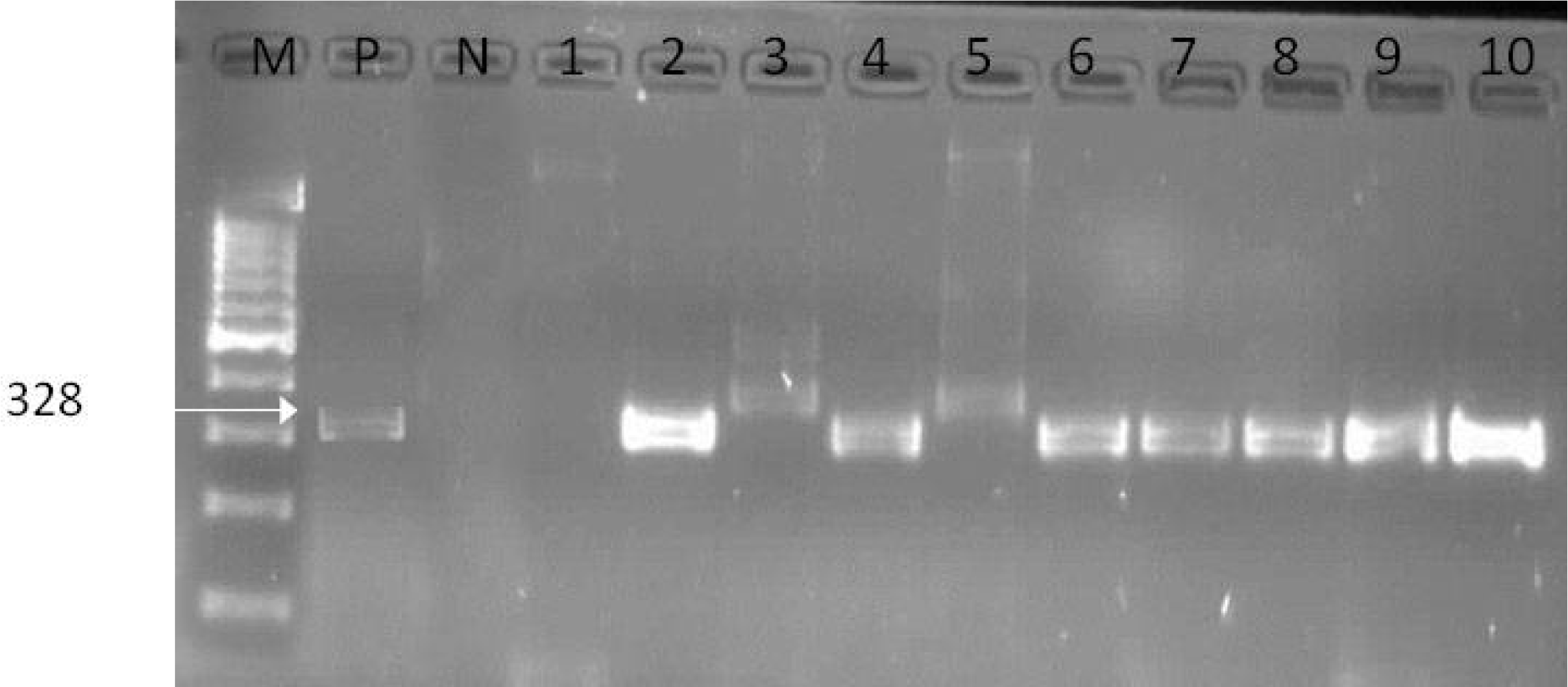
Agarose gel electrophoresis of pap gene amplification products of UPEC. M: Molecular weight marker (100bp), P: Escherichia coli (UPEC) DSM 4618 (Positive control) N: Negative control; Lanes 1-10: *E. coli* isolates

**Figure 3:**
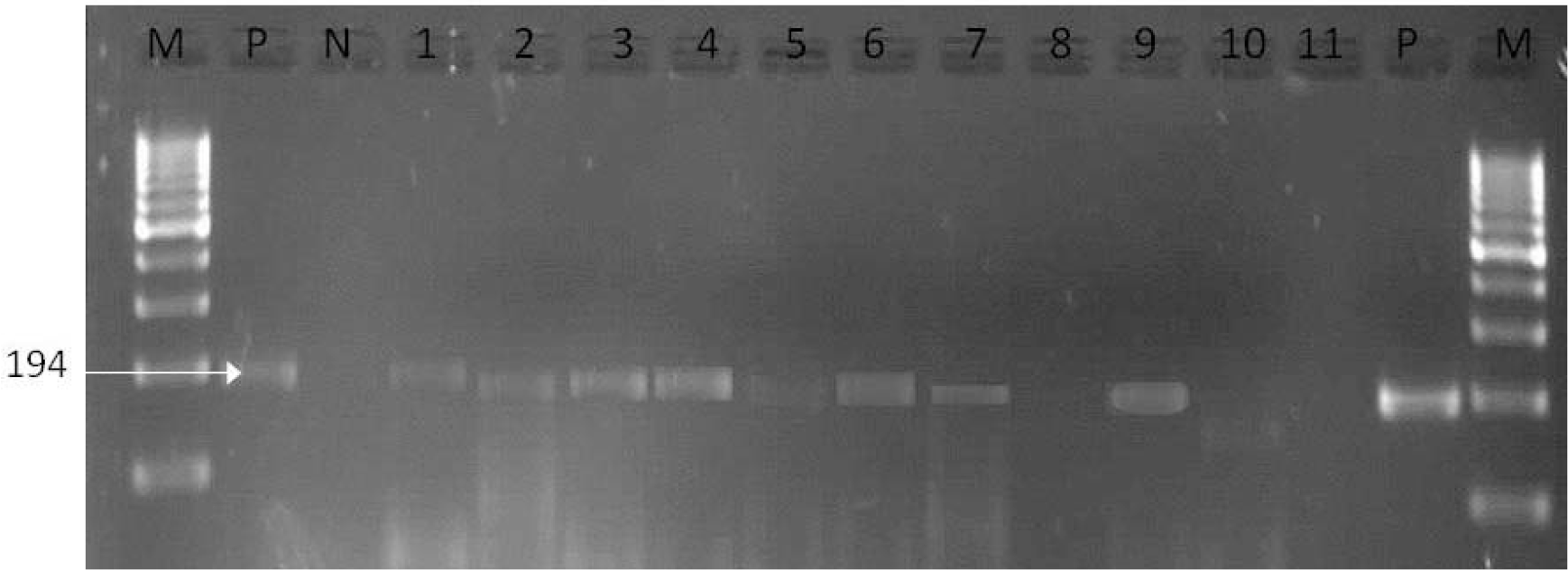
Agarose gel electrophoresis of EAgg gene amplification products of EAEC. M: Molecular weight marker (100bp), P: *Escherichia coli* (EAEC) DSM 10974 (Positive control), N: Negative control; Lanes 1-11: *E. coli* isolates

**Figure 4:**
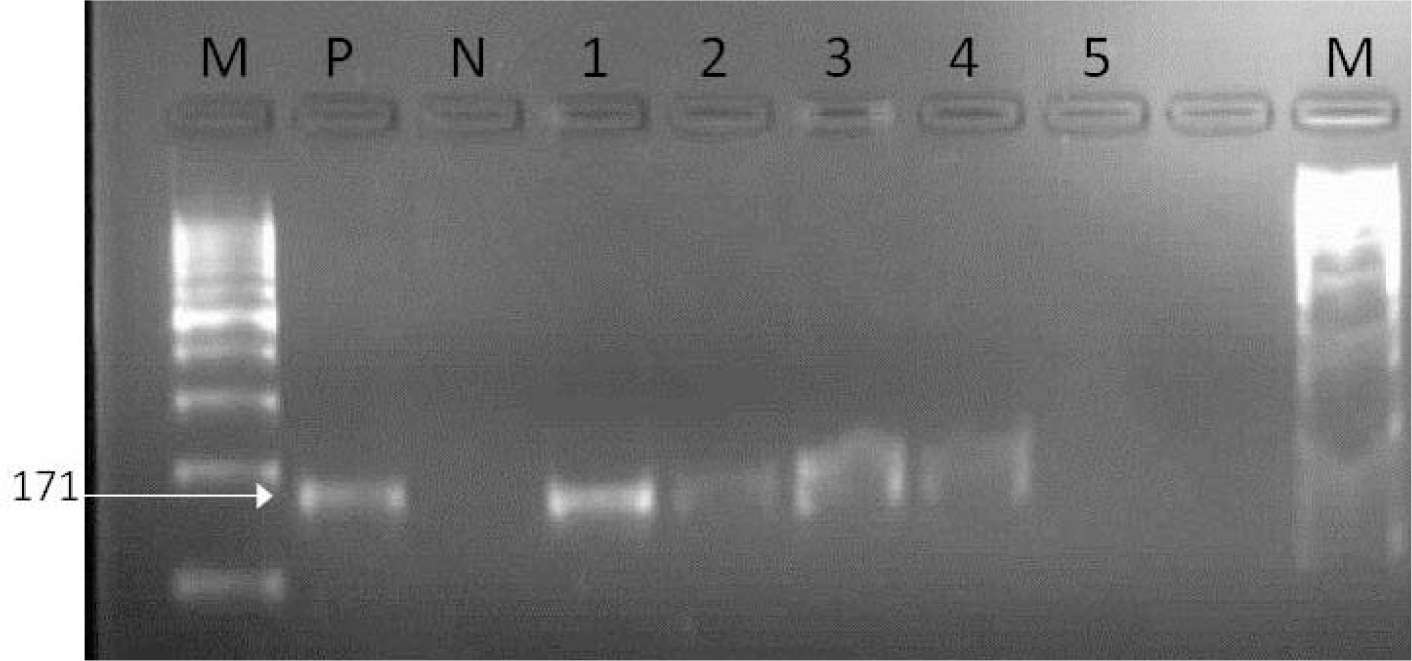
Agarose gel electrophoresis of ibe gene amplification products of NMEC. M: Molecular weight marker (100bp), P: *Escherichia coli* (NMEC) DSM 10819 (Positive control), N: Negative control; Lanes 1-5: *E. coli* isolates

### Antimicrobial susceptibility testing

The UPEC isolates showed low resistance to antibiotics like meropenem (100%), cefotaxime (100%) and gentamicin (88.9%). The isolates showed high degrees of resistance to tetracycline (74.1%), ampicillin (74.1%) and cephalothin (66.7%). Other *E. coli* pathotypes, EAEC, NMEC and EPEC, showed high sensitivity (100%) to meropenem, gentamicin and cefotaxime. EAEC had 63.6% resistance to tetracycline and 54.5% resistance to both ampicillin and cephalothin. Intermediate sensitivity (80%) to cephalothin was recorded for NMEC, which also had 60% resistance to tetracycline and 40% resistance to ampicillin. EPEC had 100% resistance to both ampicillin and cephalothin and 66.7% resistance to tetracycline. Each of the tested pathotypes showed resistance to two or three antibiotics, mainly ampicillin, tetracycline and cephalothin.

## Discussion

This study showed finding on the incidence of four pathogenic *E. coli* strains in the final effluent discharge into surface water. About eight pathogenic *E. coli* pathotypes were identified. Five of the pathotypes can cause invasive intestinal infections, watery diarrhoea and dysentery in humans and animals, while the remaining three cause extra-intestinal infections caused by extra-intestinal pathogenic *E. coli* (ExPEC) (Bekal *et al*., 2003). Four out of eight pathotypes identified and tested in this study are shown in Table 3. Both invasive and extra-intestinal pathotypes were identified. A previous study by Osode (2010) identified two *E. coli* pathotypes at Plant B; EHEC and EAEC were identified while EIEC was confirmed from another treatment Plant. This was in contrast to the outcome of this study for Plant B, in which EPEC, UPEC and EAEC pathotypes were identified. However, the detection rate at Plant B was low as compared to the second Plant (Plant A) with the exception of EPEC, which was only detected in Plant B. The efficiency of the treatment Plant could be one of the reasons why little or no pathogenic *E. coli* were detected at Plant B. Alternatively, it could be that *E. coli* strains found in these sites did not carry any virulence genes. This is evidenced by the absence of *E. coli* pathotypes from the *E. coli* that were isolated and confirmed. A similar situation was reported by Masters *et al*. (2011). However, at the Plant A, the three pathotypes were identified and in higher concentrations than Plant B. The identified pathotypes (Table 3) are of great public health importance. Apart from the EAEC previously identified by Osode (2010) in WWTP effluent in the Eastern Cape that was also identified in this study, NMEC and UPEC make up the major findings at the Plant A. Of the 476 confirmed *E. coli* isolates tested, UPEC was about 5.7% followed by EAEC at 2.3%, NMEC at 1.1% and EPEC at 0.6%. In a similar study by Verma, Ramteke and Garg (2008) in India, they reported a high incidence of UPEC in the treated final effluent as well as EPEC but at a lower concentration. Anastasi et al. (2012, 2010) demonstrated that some *E. coli* strains with uropathogenic properties survived treatment stages of sewage treatment plants and are released into the environment. The presence of EPEC in another study was found to be more common in city wastewater contrasted with slaughterhouse wastewater where the frequency of ExPEC was not influenced by the wastewater treatment process and the predominance of a characteristic EPEC was observed to be low in the final effluents (Diallo *et al*., 2013). The occurrence of EAEC in water was reported by Masters *et al*., (2011). They investigated the presence of the virulence genes attributed to EAEC. This strain was identified in conjunction with EPEC, pointing to a possible source of faecal contamination. Hamelin *et al*., (2007) reported the presence of EAEC, EPEC, UPEC and NMEC in river water receiving urban municipal wastewater. Also Koba (2013), in a study of the water from two rivers in the Eastern Cape, identified the presence of ETEC, EIEC and EPEC in one of the studied rivers and EAEC in both of the rivers studied. One of the studied sites, Plant A, also demonstrated a large diversity of *E. coli* pathotypes and similar study done by Adefisoye and Okoh, 2016, exhibit closely related trends in the quantity and types of pathogenic *E. coli* detected. The presence of these pathogenic organism groups has additionally been seen in past investigations where these strains were related with both human and non-human extra-intestinal infections (Bekal *et al*., 2003). Agricultural products and other aquaculture have been reported to have a high risk of diarrhoea as well as individuals who were in direct contact with wastewater had a higher vulnerability of acquiring disease than the individuals who were most certainly not (Trang *et al*., 2007). In the Eastern Cape and Limpopo Provinces of South Africa, these pathogenic *E. coli* with the exception of NMEC and UPEC have been isolated from diarrhoea patients, with EAEC being the predominant cause of infection (Bisi-Johnson *et al*., 2011; Samie *et al*., 2007). Their presence in the environment calls for concern because of their public health consequences (Clements *et al*., 2012)

For routine reporting and primary testing, the choices of antibiotic panels selected were based upon the recommendation of CLSI (CLSI, 2012b). The antibiotics used for this study were representatives of some different classes of antibiotic. Five classes of the antibiotics were tested and they were: ampicillin of the penicillin class, gentamicin of the aminoglycosides, tetracycline, meropenem of the carbapenems, cephalothin and cefotaxime of the first and third generation cephalosporins (CLSI, 2012b). Antibiotic profiles of the pathogenic pathotypes demonstrated the lower effectiveness of ampicillin, tetracycline, and cephalothin, which is a first-generation cephalosporin. These antibiotics constitute the major classes of antibiotic drugs commonly used in first-line treatment. Though our study never tested for other class members of tetracycline, susceptibility of organisms to doxycycline and minocycline can be considered based on their susceptibility to tetracycline. However, intermediate or resistant to tetracycline by some organisms may be susceptible to doxycycline, minocycline, or both (CLSI, 2012a). All the pathogenic isolates showed a higher level of resistance than susceptibility to tetracycline. On the average, the organisms showed 60% resistance.

The choice of cefotaxime for this study was to identify the presence of Extended-spectrum beta-lactamase (ESBL) (CLSI, 2012b) among the isolates and none were found, as demonstrated by the 100% susceptibility to the drug (Table 4). The advent of carbapenem-resistant *E. coli* has become a global concern (Nordmann *et al*., 2012), being one of the last lines defense drug for treatment. Our study was able to show that none of the pathogenic *E. coli* are resistant to the carbapenem drug class (Meropenem, see Table 4). Nontongana et al., 2014 were able to demonstrate resistance to some of these antibiotics in their study on river water in the Eastern Cape. In a study carried out in Durban on wastewater treatment plant, the *E. coli* isolates tested, the most resistance was to ampicillin, amoxicillin, doxycycline, and tetracycline (Pillay and Olaniran, 2016). Multiple resistance patterns reported by Kinge *et al*., 2010 and Mulamattathil *et al*., 2014 from wastewater, surface water and water treatment plants were similar to the resistance pattern observed for our study against ampicillin and tetracycline. Though our study didn’t find any carbapenem-resistant enterobacteriaceae (CRE) especially for *E. coli* in South Africa, but there have been reported cases of other members of the enterobacteriaceae like *Klebsiella* resistant to carbapenem (Brink *et al*., 2012). Recent reports have highlighted the need for institutions to stem the indiscriminate use of antibiotics in the country and provide restrictive measures which can curtail the looming danger of acquiring CRE in South Africa (Coetzee and Brink, 2011). The presence of antibiotics in surface water and wastewater have been identified and data collected by Matongo *et al*., 2015 reported that while insufficiently treated wastewater contributes to surface water contamination, other human activities through improper use and disposal of pharmaceutical products and wastes also contribute appreciably to the pharmaceutical loading of rivers.

**Table 4:** Antimicrobial susceptibility testing of *E. coli* pathotypes.

| Pathotypes | n = 46, Susceptibility profile (%) |  |  |  |  |  |  |  |  |  |  |  |  |  |  |  |  |  |
| --- | --- | --- | --- | --- | --- | --- | --- | --- | --- | --- | --- | --- | --- | --- | --- | --- | --- | --- |
|  | Meropenem |  |  | Gentamicin |  |  | Cefotaxime |  |  | Ampicillin |  |  | Tetracycline |  |  | Cephalothin |  |  |
|  | S | I | R | S | I | R | S | I | R | S | I | R | S | I | R | S | I | R |
| UPEC | 100 | - | - | 88.9 | 7.4 | 3.7 | 100 | - | - | 14.1 | 11.1 | 74.1 | 25.9 | - | 74.1 | 3.7 | 29.6 | 66.7 |
| EAEC | 100 | - | - | 100 | - | - | 100 | - | - | 36.4 | - | 63.6 | 36.4 | - | 63.6 | - | 45.5 | 54.5 |
| NMEC | 100 | - | - | 100 | - | - | 100 | - | - | 60 | - | 40 | 40 | - | 60 | 20 | 80 | - |
| EPEC | 100 | - | - | 100 | - | - | 100 | - | - | - | - | 100 | 33.3 | - | 66.7 | - | - | 100 |

Note: - S (susceptibility), I (Intermediate), R (Resistance)

Knowledge of the resistance pattern of pathogenic bacterial strains in geographical areas of South Africa should be enough to get their drug management and policy in place to stem the rising cases of microbial resistances in the country. It will help in directing the proper and the prudent utilization of antibiotics. The formulation of an appropriate institutional and organizational antibiotics policy will go a long way in controlling these infections (Shariff *et al*., 2013).

We have previously reported the operational status of these wastewater treatment plants often result in the discharge of inadequately treated effluent into receiving surface waters (Osuolale and Okoh, 2015b, 2015a, 2017). Time is racing for South Africa to address her water challenges. The world is calling for safe wastewater management and reuse, which formed the basis of the UN’s World Water Day. The antibiotic stewardship is an advocacy for wise antibiotic management use. It is therefore important for the management system of the Department of Water Affair to review their handling of wastewater and antibiotics wastes to minimize their environmental impacts, and public health concerns.

